## Supplementary material for "A ZTF-7/RPS-2 complex mediates the cold-warm response in *C. elegans*": Table S1 and Table S2 20220927

**Table S1: Strains used in the work.**

| Stain | Genotype |
|---|---|
| N2 | |
| SHG2103 | *3xflag::gfp::exos-1(ustIS112);rrp-8::mcherry(ustIS128)* |
| SHG2104 | *3xflag::gfp::exos-2(ustIS113);rrp-8::mcherry(ustIS128)* |
| SHG1690 | *3xflag::gfp::exos-10(ustIS114);rrp-8::mcherry(ustIS128)* |
| SHG1702 | *mCherry::dis-3(ustIS115);gfp::rrp-8(ustIS76)* |
| SHG680 | *3xflag::gfp::exos-1(ustIS112)* |
| SHG679 | *3xflag::gfp::exos-2(ustIS113)* |
| SHG1044 | *3xflag::gfp::exos-10(ustIS114)* |
| SHG1092 | *rrp-8::mCherry(ustIS128)* |
| SHG1660 | *rbd-1::mCherry(ustIS207)* |
| SHG904 | *nucl-1::gfp::3xflag(ustIS279)* |
| SHG893 | *3xflag::gfp::c27f2.4(ustIS97)* |
| SHG1257 | *3xflag::gfp::rpoa-2(ustIS116);fib-1::mCherry(ustIS140)* |
| SHG2271 | *asp-17(ust190);3xflag::gfp::exos-10(ustIS114)* |
| SHG2272 | *zip-10(ust192);3xflag::gfp::exos-10(ustIS114)* |
| SHG2085 | *glr-3(ust166);3xflag::gfp::exos-10(ustIS114)* |
| SHG1295 | *ztf-7(ust117);3xflag::gfp::exos-10(ustIS114)* |
| SHG1297 | *ztf-7(ust118);3xflag::gfp::exos-10(ustIS114)* |
| SHG1299 | *ztf-7(ust119);3xflag::gfp::exos-10(ustIS114)* |
| SHG1443 | *ztf-7(ust119);3xflag::gfp::exos-1(ustIS112)* |
| SHG1444 | *ztf-7(ust119);3xflag::gfp::exos-2(ustIS113)* |
| SHG1296 | *ztf-7(ust118)* |
| SHG1298 | *ztf-7(ust119)* |
| SHG2270 | *ztf-7(ust117);mCherry::ztf-7(ustIS115);3xflag::gfp::exos-10(ustIS114)* |
| SHG1445 | *ztf-7(ust118);mCherry::ztf-7(ustIS155)* |
| SHG1446 | *ztf-7(ust119);mCherry::ztf-7(ustIS155)* |
| YY178 | *eri-1(mg366);3xflag::gfp::nrde-3(ggIS1)* |
| SHG2182 | *ztf-7(ust118);eri-1(mg366);3xflag::gfp::nrde-3(ggIS1)* |

| | |
|---|---|
| SHG2183 | *ztf-7(ust119);eri-1(mg366);3xflag::gfp::nrde-3(ggIS1)* |
| SHG1441 | *ztf-7::gfp::3xflag(ustIS174)* |
| SHG2088 | *3xHA::rps-2(ustIS327)* |
| SHG2089 | *3xHA::rps-2(ustIS327);ztf-7::gfp::3xflag(ustIS174)* |
| SHG1560 | *ztf-7::gfp::3xflag(ustIS174);lmn-1::mCherry(ustIS144)* |
| SHG1886 | *3xflag::gfp::exos-10(D303N;E305Q)(ustIS251)* |
| SHG1887 | *3xflag::gfp::exos-10(EXO domain deletion)(ustIS252)* |
| SHG1888 | *3xflag::gfp::exos-10(HRDC domain deletion)(ustIS253)* |
| SHG2267 | *ztf-7(ust119);3xflag::gfp::exos-10(D303N;E305Q)(ustIS251)* |
| SHG2268 | *ztf-7(ust119);3xflag::gfp::exos-10(EXO domain deletion)(ustIS252)* |
| SHG2269 | *ztf-7(ust119);3xflag::gfp::exos-10(HRDC domain deletion)(ustIS253)* |

**Table S2. Sequences of sgRNAs for CRISPR/Cas9-mediated gene editing.**

| | |
|---|---|
| *ztf-7::gfp* sgRNA#1 | TTCGTTATACATAGTACATTCGG |
| *ztf-7::gfp* sgRNA#2 | ATGTACTATGTATAACGAATAGG |
| *ztf-7::gfp* sgRNA#3 | ACAACAATCTAATGATTGATAGG |
| *mCherry::ztf-7* sgRNA#1 | GAAATCGCCGACTTGCGAGGAGG |
| *mCherry::ztf-7* sgRNA#2 | GCAATGACTAACCGATTTTCGGG |
| *mCherry::ztf-7* sgRNA#3 | TTCGGGATAATTGAGATGAGGGG |
| *ztf-7* sgRNA#1 | ATCAATCCAATCGTAAGGACTGG |
| *ztf-7* sgRNA#2 | ATGAGCTTGACTGTTGCTCGCGG |
| *ztf-7* sgRNA#3 | AAGTAGGAATCGCCTTCGCTTGG |
| *glr-3* sgRNA#1 | TCTTGCTTGATCCTCCACAATGG |
| *glr-3* sgRNA#2 | GACAGGTGAAGACTCACACATGG |
| *glr-3* sgRNA#3 | CAATTGGTTCGACTAAAGCATGG |
| *asp-17* sgRNA#1 | CTGCAGAGCAGAAAGGAGCTAGG |
| *asp-17* sgRNA#2 | GGGCAGCATGCTGAGTAATCAGG |
| *asp-17* sgRNA#3 | TGTTGGCAGATCCAGTGTCAAGG |
| *zip-10* sgRNA#1 | GCTGATGAGAATAGAGATTCAGG |
| *zip-10* sgRNA#2 | CTGAATTGAGCAAGCATTGCTGG |
| *zip-10* sgRNA#3 | AACATCTACCACATCCAGCTCGG |
